# Endogenous Galectin-3 is Required for Skeletal Muscle Repair

**DOI:** 10.1101/2020.10.01.322461

**Authors:** Daniel Giuliano Cerri, Lilian Cataldi Rodrigues, Vani Maria Alves, Juliano Machado, Víctor Alexandre Félix Bastos, Isis do Carmo Kettelhut, Luciane Carla Alberici, Sean R. Stowell, Maria Cristina R. Costa, Richard D. Cummings, Marcelo Dias-Baruffi

**Affiliations:** Department of Clinical Analyses, Toxicology and Food Sciences, School of Pharmaceutical Sciences of Ribeirão Preto, University of São Paulo, Ribeirão Preto, SP, Brazil; Department of Cellular and Molecular Biology and Pathogenic Bioagents, School of Medicine of Ribeirão Preto, University of São Paulo, Ribeirão Preto, SP, Brazil; Department of Physiology, Ribeirão Preto Medical School/University of São Paulo, Ribeirão Preto, SP, Brazil; Department of Biochemistry/Immunology, Ribeirão Preto Medical School/University of São Paulo, Ribeirão Preto, SP, Brazil; Department of BioMolecular Sciences, School of Pharmaceutical Sciences of Ribeirão Preto, University of São Paulo, Ribeirão Preto, SP, Brazil; Pathology Department, Emory University School of Medicine, Atlanta, Georgia, USA; Estácio University Center- Ribeirão Preto, SP, Brazil; Department of Surgery, Beth Israel Deaconess Medical Center, Harvard Medical School, 3 Blackfan Circle, Room 11087, Boston, MA, 02115, USA

**Author notes:** Corresponding author at: Department of Clinical Analyses, Toxicology and Food Sciences, School of Pharmaceutical Sciences of Ribeirão Preto, University of São Paulo, Ribeirão Preto, SP, Brazil. E-mail address (M. Dias-Baruffi). **Abbreviations:** Akt= serine/threonine protein kinase; also called protein kinase B; CREB= cAMP response element-binding protein; DM= Differentiation medium; ERK= extracellular signal–regulated kinase; Gal-3, -1, -2, -7 and 12= Galectin-3, -1, -2, -7, respectively; *GALKO-3*= Galectin-3 knockout mice; GM= Growth medium; HGF= Hepatocyte Growth Factor; IGF-1= Insulin Growth Factor-1; *mdx*= dystrophic mouse model; MMP= Matrix Metalloproteinases; MRP= Muscular Repair Process; MyoD= Myogenic differentiation factor; OPN= Osteopontin; Pax7= Paired Box 7; PBS= phosphate buffered saline; TIMPs= Tissue Inhibitor of Metalloproteinase, TBS= Tris buffered saline; WT= Wild Type mice for *GAL3KO*, wt= wild type for *mdx* mice.

**Keywords:** Galectin-3, Myolesion, Muscle repair, Myoblast, Myogenic factors, Myotubes

## Abstract

Skeletal muscle has the intrinsic ability to self-repair through a multifactorial process, but many aspects of its cellular and molecular mechanisms are not fully understood. There is increasing evidence that some members of the mammalian β-galactoside-binding protein family (galectins) are involved in the muscular repair process (MRP), including galectin-3 (Gal-3). However, there are many questions about the role of this protein on muscle self-repair. Here, we demonstrate that endogenous Gal-3 is required for: i) muscle repair *in vivo* using a chloride-barium myolesion mouse model, and ii) mouse primary myoblasts myogenic programming. Injured muscle from Gal-3 knockout mice (*GAL3KO*) showed persistent inflammation associated with compromised muscle repair and the formation of fibrotic tissue on the lesion site. In *GAL3KO* mice, osteopontin expression remained high even after 7 and 14 days of the myolesion, while MyoD and myogenin had decreased their expression. In *GAL3KO* mouse primary myoblast cell culture, Pax7 detection seems to sustain even when cells are stimulated to differentiation and MyoD expression is drastically reduced. These findings suggest that the detection and temporal expression levels of these transcriptional factors appear to be altered in Gal-3-deficient myoblast cell culture compared to Wild Type (WT) cells. We observed Gal-3 expression in WT states, both *in vivo* and *in vitro*, in sarcoplasm/cytoplasm and myonuclei; as differentiation proceeds, Gal-3 expression is drastically reduced, and its location is confined to the sarcolemma/plasma cell membrane. We also observed a change in the temporal-spatial profile of Gal-3 expression and muscle transcription factors levels during the myolesion. Overall, these results demonstrate that endogenous Gal-3 is required for the skeletal muscle repair process.

## INTRODUCTION

The adult skeletal muscle is a specialized mammalian tissue that has a notable ability to regenerate after injury. In normal conditions, few quiescent and partially differentiated satellite cells (stem cells) located among muscle fibers are fused in a turnover process responsible for daily repair of micro lesions caused by movement and force (Chargé and Rudnicki, 2004; Gharaibeh et al., 2012). This occurs due to the muscle replenishing “nursery” (Mukund and Subramaniam, 2020; Zhuang et al., 2020). In chronic or acute injured skeletal muscles, however, the molecular and cellular muscle repair processes (MRP) heavily depend on the dynamic interaction between satellite cells (SC) and its microenvironment. SC are characterized by the expression of the transcription factor Pax7 (Asfour et al., 2018; Gharaibeh et al., 2012; Hernández-Hernández et al., 2017; Hindi et al., 2013; Péault et al., 2007), and upon its activation, SC become proliferative myoblasts. Pax7 levels decrease, along with MyoD transcription factor expression, which drives myoblasts cells to proliferation and repairing process. The consequent alignment with pre-existing myofibers and subsequent fusion results in regenerated muscle fiber (Asfour et al., 2018; Chargé and Rudnicki, 2004; Ten Broek et al., 2010). Other factors, such as growth factors IGF-1, HGF, and osteopontin (Pagel et al., 2014; Tatsumi et al., 2006) are released from the damaged fiber, triggering the MRP. Several intracellular events will culminate in this cell-to-tissue reorganization such as the phosphorylation of key proteins such as Akt, p38MAPK, ERK and CREB (Hindi et al., 2013; Magenta et al., 2003; Segalés et al., 2016; Stewart et al., 2011) Pro-inflammatory mediators, such as IL-6, IL-1, and TNF-α, which are secreted by SC and contribute to immune cell infiltration by M2 macrophages, which along with other cells in the muscular injury, induce myogenesis (Cantini et al., 2002; Mantovani et al., 2005; Tidball, 1995).

In addition to the cells and mediators already described, other soluble proteins from the lectins family seem to contribute to MRP. Galectins are multifunctional β-galactoside-binding lectins, expressed in various tissues and in immunological cells, such as monocytes and macrophages, playing important roles in biological events, both in acute and chronic inflammation (Cerliani et al., 2011; Liu and Rabinovich, 2010). The cell-cell and cell-extracellular matrix interaction by galectins may be responsible for cross-links events between receptors triggering cell membrane alterations, signal transduction, migration, and others, such as membrane fusion, as occur in the myogenesis process (Barondes et al., 1994). The 14 kDa lectin, Galectin-1 (Gal-1), was the first and most studied galectin in MRP (Goldring et al., 2002). It was discovered as a primordial factor in the process of muscle differentiation, transdifferentiation of mesenchymal cells to myogenic lineage and in myogenesis (Ahmed et al., 2009; Cerri et al., 2008; Ding et al., 2016; Shoji et al., 2009). The presence of Gal-1 colocalized with CD45^+^ cells infiltrating leukocytes in injured tissue demonstrates this lectin participation on muscle degeneration followed by repair (Cerri et al., 2008). Besides Gal-1, a broad spectrum of lectins has emerged playing important roles during the myogenic programming process. During myoblast differentiation to myotubes, in C2C12 cell lineage, Gal-2 mRNAs are downregulated, and mRNAs from Gal-7 and Gal-12 are substantially upregulated (Janot et al., 2009).

Recently, Gal-3 was observed to increase the efficiency of myogenesis in a C2C12 cell lineage, being expressed in both myoblasts and myotubes during differentiation (Rancourt et al., 2018). Gal-3 is a chimeric lectin (∼30 kDa monomer) which contains two domains: (i) C-terminal carbohydrate recognition domain and (ii) an N-terminal collagen-like rich in proline and glycine residues that promotes formation of functional oligomers (Varki et al., 2017). Interestingly, Gal-3 can be found in the nucleus, cytoplasm, cell surface and as a secreted protein (Arthur et al., 2015a; Haudek et al., 2010; Mehul and Hughes, 1997). As a multifunctional lectin, Gal-3 can bind glycoproteins on cell surfaces and within the matrix, including integrins, fibronectin, and collagen, triggering effector functions in cell migration, apoptosis, fibrosis, and inflammation (Dumic et al., 2006; Ochieng et al., 2002; Sato et al., 2009). Although Gal-3 is expressed in all types of immune cells, endothelial cells and others, it is also expressed in skeletal, smooth and cardiac muscle, being down or up-regulated according to the inflammatory cellular microenvironment (Dong et al., 2018). For example, in cardiac disease models, the elevated levels of Gal-3 identify it as a biomarker for myocardial fibrosis, linked with Damage-Associated Molecular Patterns (DAMPs) (Henderson et al., 2006; Lok et al., 2010; Sato et al., 2009). In this context, macrophage-derived Gal-3 induces cardiac fibroblast proliferation and collagen deposition (Sharma et al., 2004).

In the present study, we investigated whether the absence of Gal-3 could restrict the MRP after barium-induced acute myolesion *in vivo* and in the myogenic programming process in mouse primary myoblasts cells *in vitro*. Also, we evaluated the location of Gal-3 during the myoblasts differentiation process and its effect on the expression of muscle transcription factors. Our results indicate that Gal-3 plays a key role in the repair of muscle cell.

## MATERIAL AND METHODS

### Animals

Adult C57BL/6 (WT) and Gal-3 knockout (*GAL3KO*) male mice (7–10 weeks old); C57BL/10 (wt) and *mdx* (for dystrophic model-4-10 weeks old) were housed at the animal facility of the Faculty of Pharmaceutical Sciences of Ribeirão Preto University of São Paulo, Brazil. The animal handling was in accordance with the guidelines set by the Institutional Laboratory Animal Care Committee of the University of São Paulo (process no. 2014.1.478.53.4).

### Antibodies and reagents

The polyclonal chicken-derived anti-Gal-3 antibody was produced in our laboratory. The human recombinant Gal-3 (hr-Gal-3) protein was prepared as described elsewhere (Arthur et al., 2015b), antibodies anti-dystrophin (sc-153760), anti-desmin (sc-7559), anti-myogenin (sc-576) and anti-pCREB (sc-9198) were purchased from Santa Cruz Biotechnology (Santa Cruz, CA); anti-pAkt (#9271) and anti-β-tubulin (#2146) from Cell signaling; anti-β1-Integrin (CSAT) and anti-osteopontin from DSHB (Developmental Studies Hybridoma Bank at University of Iowa); anti-MyoD (ThermoFisher, clone 5.8A); Anti-Pax7 (Millipore MAB020 2F-12H4); donkey anti-rabbit-Alexa_546, donkey anti-mouse-Alexa_546, Goat anti-Chicken-Alexa_488 from Molecular Probes® Thermo Fisher Scientific; donkey IgG, donkey anti-goat HRP, donkey anti-mouse HRP, donkey anti-rabbit HRP and Donkey anti-chicken HRP from Jackson Immunoresearch (West Groove, PA). DAPI (4’,6-Diamidino-2-Phenylindole, Dihydrochloride) and Hoechst 33342, Trihydrochloride, Trihydrate from Thermo Scientific.

### BaCl_2_-induced lesion

Adult C57BL/10 and *GAL3KO* male mice (7–10 weeks old) were anesthetized intraperitoneally with 100 mg/kg sodium thiopental combined with 10 mg/Kg lidocaine. Subsequently, 50 μL of 1.2% sterile barium chloride solution (BaCl_2_, Carlo Erba, Milan, Italy) was injected into the gastrocnemius muscle.

### Immunofluorescence in muscle tissue

Immunofluorescence assay was performed as described in (Cerri et al., 2015). Briefly, frozen 5 μm tissue sections were blocked in TBS with 50 μg/mL donkey IgG, 2% (w/v) BSA, and 0.5% (v/v) Triton X-100 for 45 min, and then incubated with 10 μg/mL chicken anti-Gal-3, 10 μg/mL anti-dystrophin, 1:25 anti β1-integrin and 1:50 anti-osteopontin for 1 h. Slides were washed twice for 15 min in Tris buffered saline (TBS) plus 0.1% (v/v) Triton X-100® and twice in TBS plus 0.05% (v/v) Tween 20®. Subsequently, slides were incubated with 5.7 μg/mL of the correspondent secondary antibody plus 50 ng/mL DAPI (4,6-diamidino-2-phenylindole, dihydrochloride) for 1 h. The washing procedure was repeated using only TBS plus Tween 20®. Slides were briefly washed in dH_2_O and finally mounted in Fluoromount: TBS (2:1). Images were captured using a Leica TCS-SP5 confocal microscopy (Mannheim, Germany).

### Histopathological analyses by Hematoxylin/Eosin (HE), Sirius Red and Masson’s Trichrome

Gastrocnemius muscle from WT and *GAL3KO* animals were fixed in 4% paraformaldehyde overnight at 4°C and paraffin-embedded for histopathological analysis. Briefly, 6 μm of tissue sections were deparaffinized and rehydrated according to standard protocols. All samples were stained routinely with Hematoxylin-Eosin (HE), Sirius Red and Masson’s Trichrome, at room temperature, as described in the sub-items below. Following the specific coloration, tissue sections were dehydrated and mounted with mounting medium (Tissue-Tek, Sakura, Japan). Analyses were performed under a light microscope Nikon Eclipse 800 with a Nikon DMX 1200 camera.

### Hematoxylin-Eosin staining

First, the rehydrated tissues were stained with Harris’ hematoxylin solution (16.5 mM Hematoxylin, 194 mM potassium aluminum sulfate and 11.54 mM mercuric II oxide, red) for 1 min and then rinsed in tap water until the water was colorless. Next, they were washed three times in dH_2_O. The following procedure was incubation in eosin solution (1.60 mM Eosin Y, 0.13 mM Floxin B, 0.45% acetic acid in final Ethanol concentration of 83%) for 30 seconds and then incubation in acid solution of ethanol 95% for 2 min.

### Sirius Red staining

Rehydrated tissues were incubated with Sirius Red solution (0.73 mM Sirius Red in saturated picric acid) for 20 seconds and then rinsed in tap water. Next, the sections were incubated in Harris’ hematoxylin solution for 1 min. They were washed twice in tap water.

### Masson’s Trichrome

The first incubation was done in Harris’ hematoxylin solution for 1 min, followed by 3 min in tap water and twice in dH_2_O for 15 seconds. Then, the sections were incubated in Ponceau acid fuchsin solution (7.4 mM Fuchsine, 3.7 mM Red Ponceau in 1% acetic acid) for 2 min, followed by washing in dH_2_O. Two other incubations were performed: 1% phosphomolybdic acid for 2 min followed by 2.5% aniline blue for 3 min. Samples were then incubated for 1 min in 1% acetic acid and in 1% acetic acid in 95% ethanol.

### Western blotting of muscle tissue

Crude muscle protein extracts were obtained by homogenization of weighted frozen tissue with a buffer solution (62.5 mM Tris, pH 6.8, 2% (w/v) SDS, 5% (v/v) glycerol, 0.10 mM SERVA blue, 0.25 mM EDTA, 30 μM phenol red, and 0.9% β-mercaptoethanol) in liquid nitrogen. The samples were incubated for 5 min at 100°C and centrifuged for 5 min at 20,000 x g. Samples were diluted at 500 μg/well and loaded into 12 or 15% polyacrylamide gels. Nitrocellulose electro blotted membranes (Amersham Biosciences, Uppsala, Sweden) were probed with 1.5 μg/mL chicken polyclonal anti-Gal-3; all commercial antibodies were used in according to manufacturers’ instructions; and then incubated with 40 ng/mL of a specific HRP conjugate secondary antibody. Bounded antibodies were revealed by enhanced chemiluminescence using the *Luminata Forte* HRP Substrate (Millipore, Billerica, MA).

### Myoblast cell culture and myotubes formation

Gastrocnemius muscles were collected and washed three times in sterile PBS. Mechanical fragmentation was done in a sterile petri dish with PBS containing 5.6 mM glucose and 1% penicillin, using a small curved *Metzenbaum* scissor and a 22-blade scalpel, both sterile. After obtaining minimal muscle fragments, type II collagenase diluted in free-serum DMEM (previously sterilized with a 0.22 μm pipette filter) was added. The sample was incubated at 5% CO_2_ and 37°C for 90 min under agitation. To neutralize the collagenase, 1 mL of fetal bovine serum was added to the suspension. Samples were centrifuged at 200 x g for 20 min at 4°C. The pellet was resuspended in 1 mL of Growth Medium-GM (DMEM high glucose supplemented with 10% fetal bovine serum, 10% horse serum, 1.5 g/L sodium bicarbonate, 0.434 mg/mL glutamine, 100 U/mL penicillin, 100 μg/mL streptomycin, and 0.25 μg/mL amphotericin B). Cells were cultured with GM at 37°C and 5% CO_2_ in a humidified cell incubator (Thermo Forma). Depending on the assay, cells were added to a six well plates or glass coverslips both previously pre-coated with Matrigel® following the manufacturer’s instructions. For myoblast differentiation to myotubes, GM was replaced for Differentiation medium-DM (replacing the serum on GM by 2% horse serum).

### Immunofluorescence in cell culture

Cells (5.0×10^4^) were cultured overnight on 13 mm round coverslips pretreated with Matrigel® as previously described. Cells were fixed for 20 min with 4% formaldehyde, 50 μM taxol (Sigma-Aldrich) and 50 mM EGTA (Sigma-Aldrich) in PBS for 20 min at 37°C. Next, cells were permeabilized with 0.3% Triton in PBS for 10 min at RT. For observation of Gal-3 on myoblast/myotubes cell surface, this last step was suppressed. The blockage of nonspecific sites was performed by 1% BSA and 5 μg/mL donkey-IgG (Jackson ImmunoResearch) in PBS for 45min. The cells were then labeled with the primary antibody diluted in PBS + 1% BSA (Sigma-Aldrich) for 1h at RT. After incubation the cells were rinsed thoroughly in PBS and the samples incubated for 45 min at RT with the secondary antibody diluted in PBS plus 0.4 μM DAPI. For nuclei staining on non-permeabilized cells, we used Hoechst 33342 at final concentration of 1μg/mL. All samples were then rinsed in PBS and once quickly in dH_2_O to mount coverslips with Fluoromount-G (EM Sciences).

### Relative quantification of Gal-3 and MyoD in immunofluorescence acquisitions

By the analysis of several micrographs, aided by the software of the confocal microscope LEICA LAS AF Lite, regions of interest with predominance of myoblasts and myotubes were separately selected and relatively quantified in terms of gray values.

### Western blotting of primary cell culture

For immunoblotting of whole cell lysates, monolayer cells (5.0×10^5^) were cultured overnight on 6 wells plates pretreated with Matrigel®. Cells were immediately lysed with a hot sample buffer (125 mM Tris-HCl, pH 6.8, 4% SDS, 10% glycerol, 1.8% 2-mercaptoethanol, 0.006% bromophenol blue). Cell lysates were boiled for 10 min and the proteins were separated by SDS-PAGE, transferred to Hybond® membranes (GE-Healthcare) and blotted with the indicated antibodies.

## RESULTS

We initiated two major experimental approaches regarding the muscle tissue: *In vivo* assessments involving myolesion induced by barium in Figures 1, 2 and 3, and *in vitro* assessments, involving primary myoblast cell culture in Figures 4, 5 and 6.

**Figure 1.**
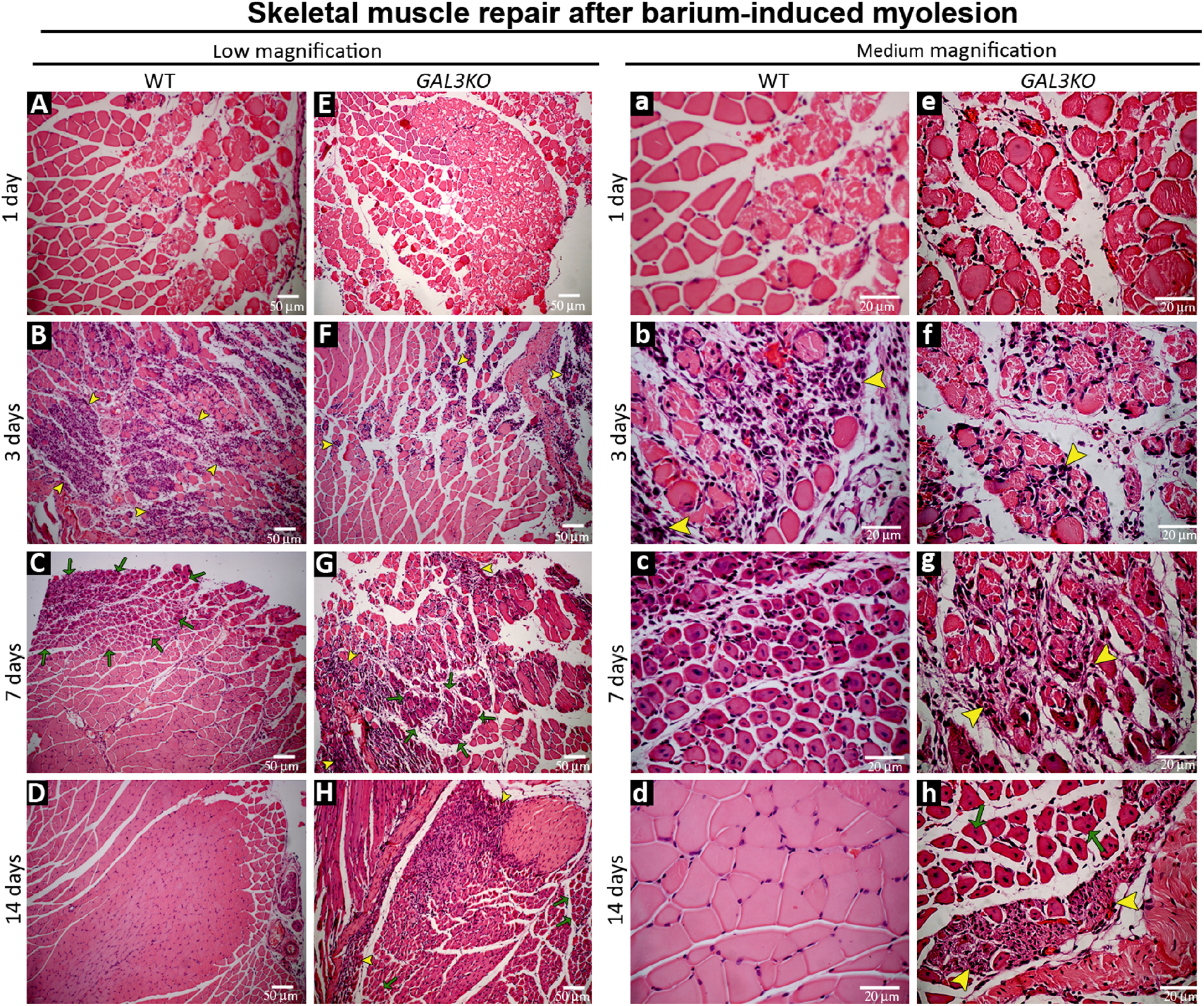
Histopathological analysis of muscle tissue repair after barium chloride-induced myolesion in WT gastrocnemius (A-D and its magnification in a-d) and *GAL3KO* (E-H and its magnification in e-h). One day post myolesion in A, E, a and e; 3 days in B, F, b and f; 7 days in C, G, c and g; 14 days in D, H, d and h. In WT muscle, regions of degenerated tissue are seen at 1 day (A and a). On the third day post myolesion (B and b), an intense inflammatory infiltrate is seen. After 7 days, areas of regenerating myofibers with non-peripheral nuclei were observed (C, green arrows and c, green arrows). Within 14 days, skeletal muscle was completely repaired (D and d). On the other hand, *GAL3KO* muscles presented a similar degenerated area after one day (E and e), but in the third day (F and f) the inflammatory infiltrate appears to be less intense than that observed on WT. Interestingly, after 7 days (G and g) the *GAL3KO* muscle presented inflammatory infiltrate (G, yellow arrowheads and g, yellow arrowheads) and some areas with regenerating myofibers (G, green arrows). Fourteen days after myolesion, the absence of Gal-3 leads to a persistent inflammatory infiltrate in the muscle (H, yellow arrowheads and h, yellow arrowheads) and regenerating myofibers (H, green arrows and h, green arrows).

**Figure 2.**
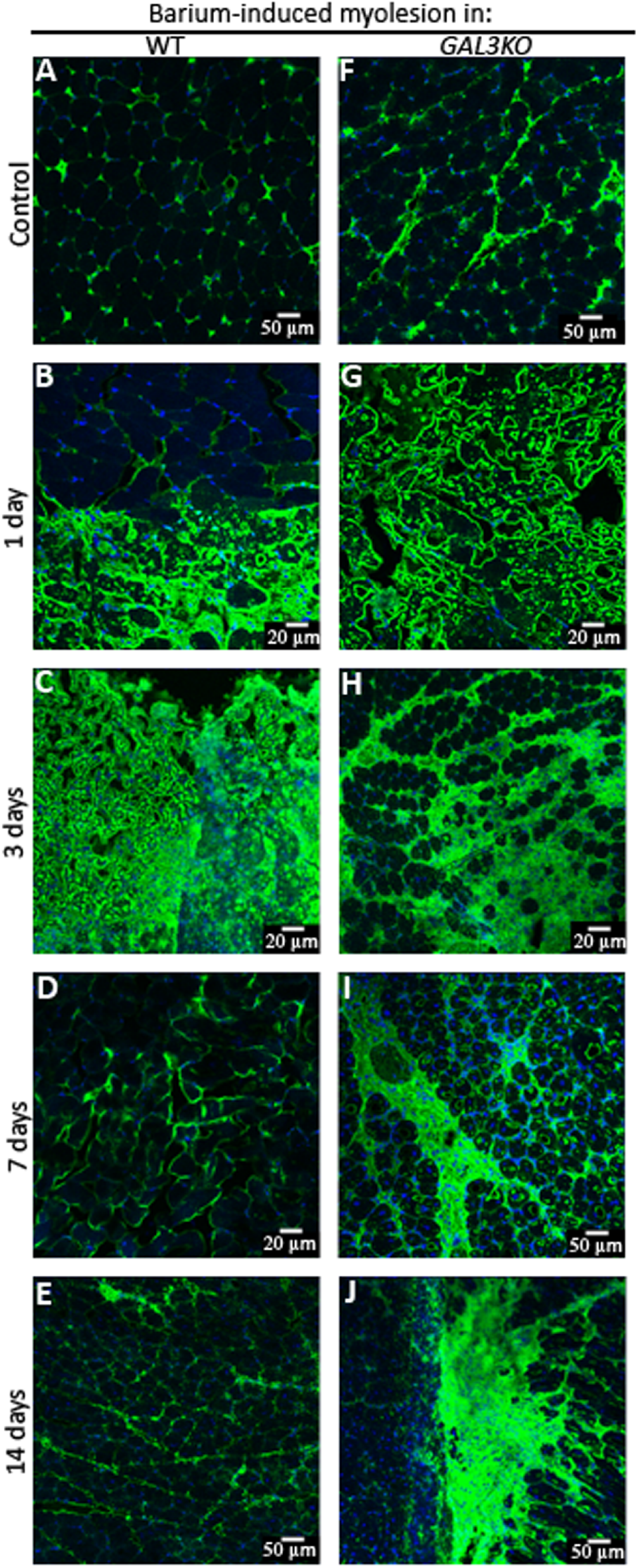
Analysis of the immunolocalization of osteopontin during the evolution of muscle tissue repair in WT (A-E) and *GAL3KO* (F-J) gastrocnemius after myolesion. An intense distribution of OPN in the injured regions after 1 and 3 days in WT animals (B and C, respectively) and *GAL3KO* (G and H, respectively) is observed. After 7 and 14 days, WT muscle (D and E, respectively) show distribution of OPN around myofibers. On the 7th and 14th days post myolesion, *GAL3KO* animals (I and J, respectively) showed an intense distribution of OPN on regions suggestive of fibrosis, besides its location around myofibers.

**Figure 3.**
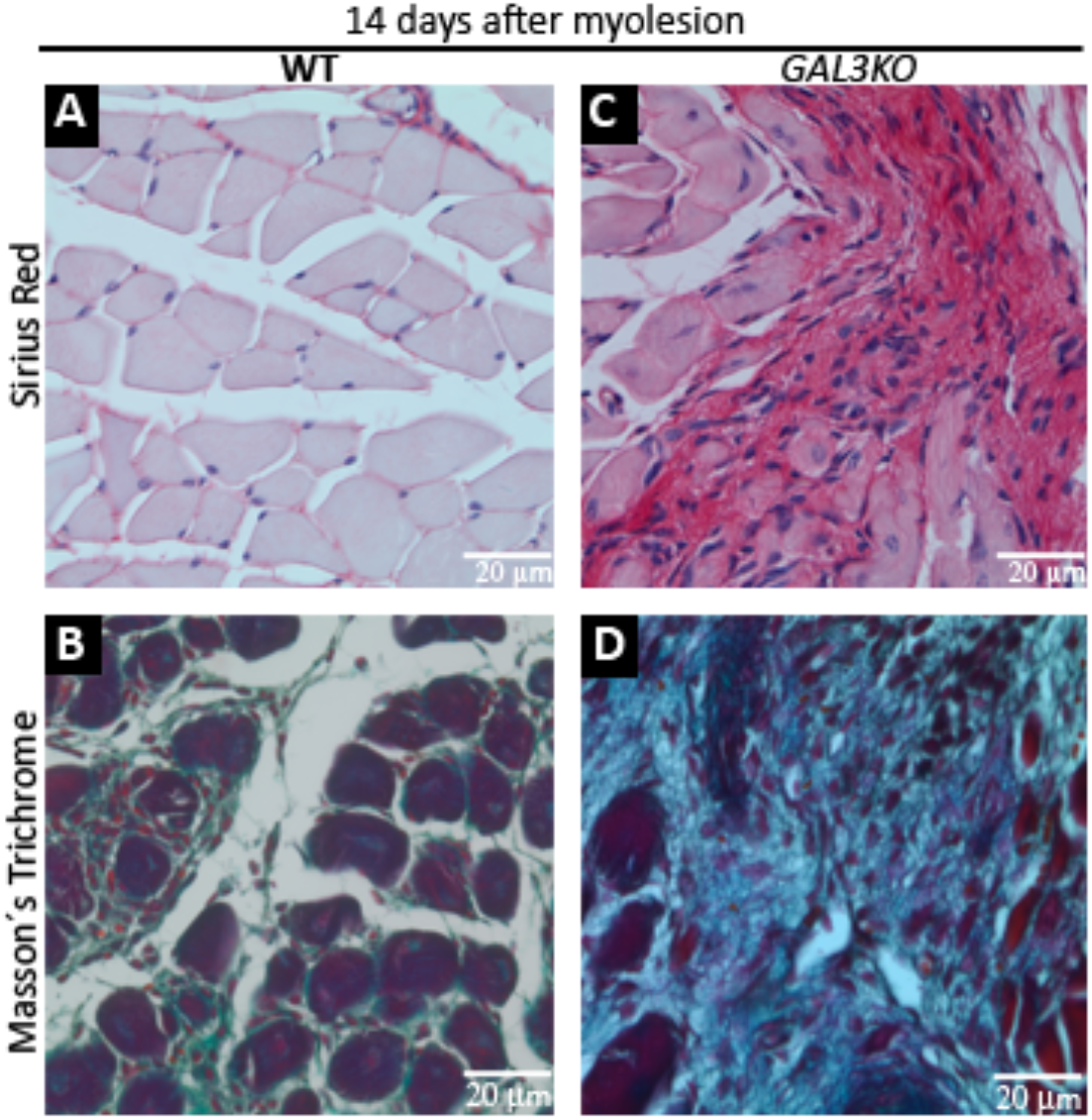
Analysis of the scar tissue formation at 14 days post-myotrauma in WT (A and B) and *GAL3KO* (C and D). To confirm scar tissue formation after 14 days of myolesion, we show here two colorations for collagen: Sirius Red (A and C) and Masson’s Trichrome (B and D). The strong reddish (C) and blueish (D) coloration indicates (both) the presence of collagen fibers in *GAL3KO* muscle.

**Figure 4.**
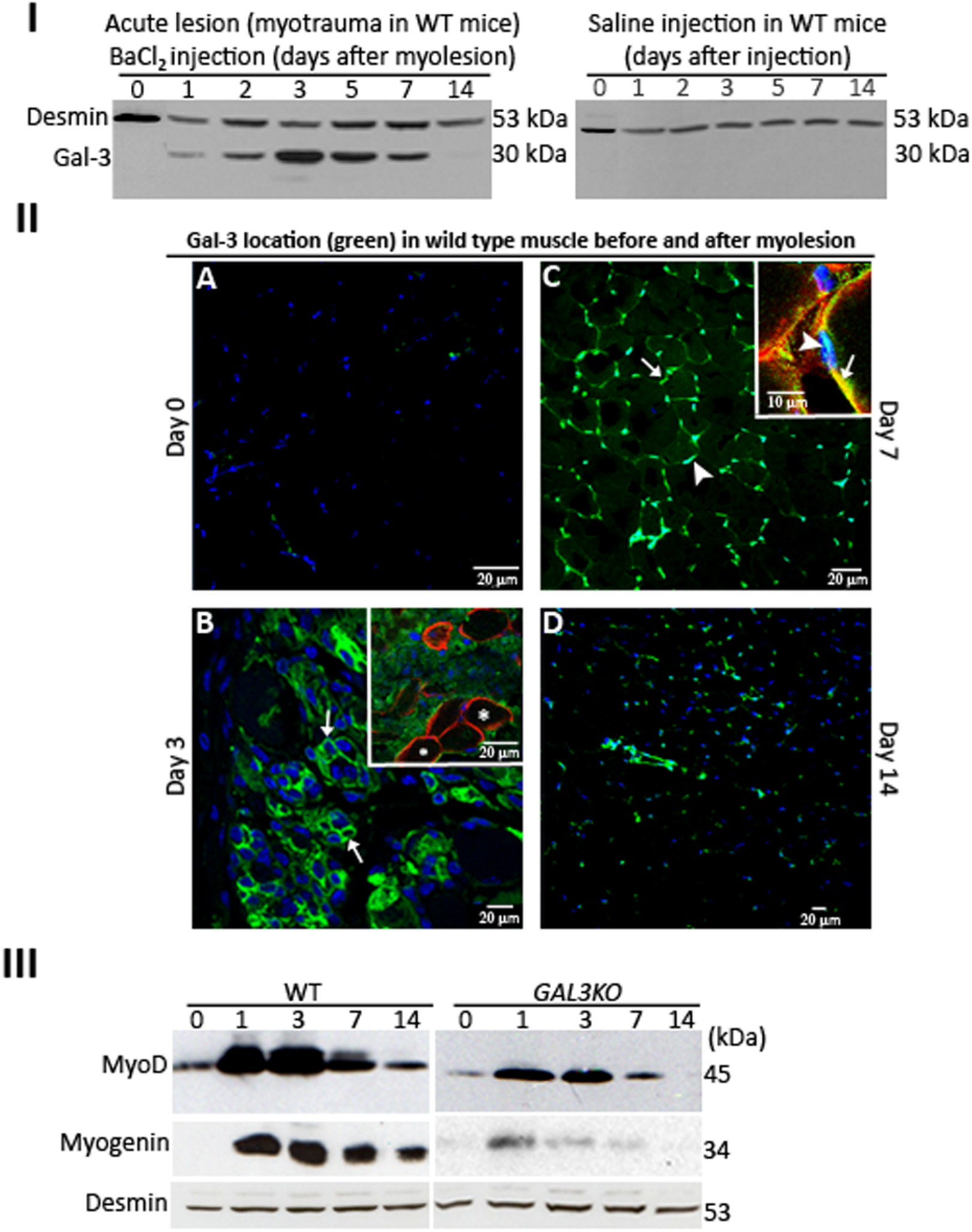
Acute myolesion induces transient Gal-3 overexpression (I) with a multitude of locations in tissue structures (II-A-D) and the absence of Gal-3 impaired MyoD and Myogenin expression (III) during acute lesion. Gal-3 was probed during the evolution of muscle tissue repair after barium chloride-induced myolesion in WT gastrocnemius (I and II). Gal-3 peaked its expression within 3 days (I-left panel), decreasing its expression as repairing process goes on. In regards of Gal-3 localization during MRP (II-A to D), in injured muscle, Gal-3 was practically undetectable (II-A), however the lectin pattern on the third day post-injury could be visualized in sarcolemma (II-B) and areas of inflammatory infiltrate (II-B, inset). Besides, Gal-3 is not detected inside the intact myofibers, as seen by dystrophin staining (II-B, inset-Asterisk). At 7 days, Gal-3 is located on myonuclei (II-C, arrowhead) and in the sarcolemma (II-C, arrow). The inset (in II-C) shows Gal-3 co-localization with both β1-integrin (II-C, inset-arrow) and DAPI (II-C, arrowhead). After 14 days (II-D), Gal-3 is detectable in a few punctuated areas. The transcription factors MyoD and Myogenin have their expressions impaired after myolesion in GAL3KO (III-right panel) when compared with WT (III-left).

**Figure 5.**
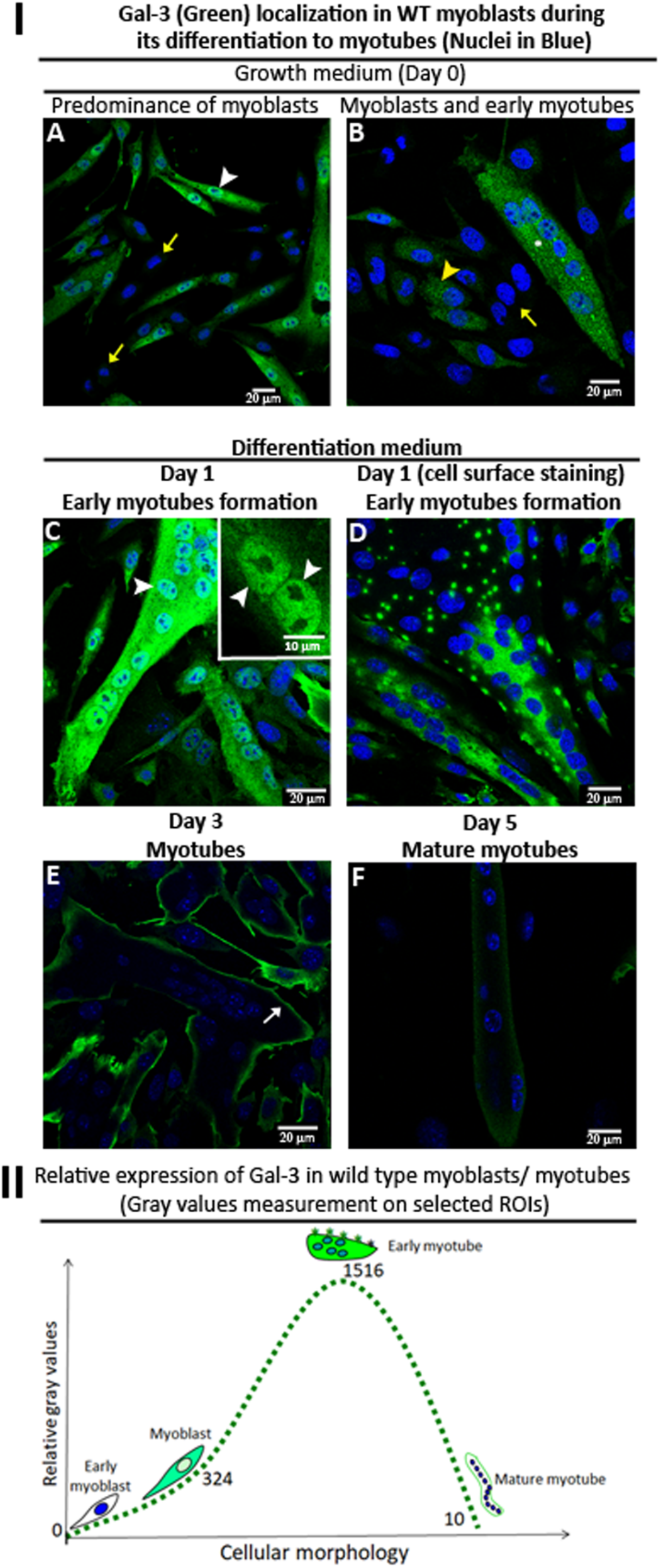
Primary myoblast cell culture from WT animals showed a multitude of Gal-3 locations during myoblast differentiation (I); Gal-3 relative expression varies abruptly as differentiation proceeds (II). Cells on growth medium (I-A and I-B) and in differentiation medium (I-C, I-D, I-E and I-F). Zero day in I-A and I-B, one day of differentiation in I-C and I-D, three days in I-E and finally, five days in I-F. Gal-3 (green) and nuclei (blue) merged localization during the dynamics of myoblast differentiation to myotubes showed a pleiotropic distribution. Gal-3 is not detected in some myoblasts (I-A and I-B, yellow arrows); its expression increases during the early cells’ alignment (I-B, yellow arrowhead); myoblast with a prominent fusiform morphology presented high Gal-3 expression (I-A, arrowhead). At the nascent myotubes (I-C) Gal-3 is located throughout its cytoplasm and in to the myonuclei (C, arrowhead and C, inset-arrowhead); outside the nascent myotubes, Gal-3 is located in several conglomerates (I-D). As differentiation proceeds, Gal-3 becomes restricted to the plasma membrane (E, arrow). After completed maturation (F), its probing is almost undetectable. The bottom scheme (II) represents the relative expression of Gal-3 during myoblast differentiation; it has a high and transient expression during the formation of nascent myotubes, returning to undetectable levels as the myotubes mature.

**Figure 6.**
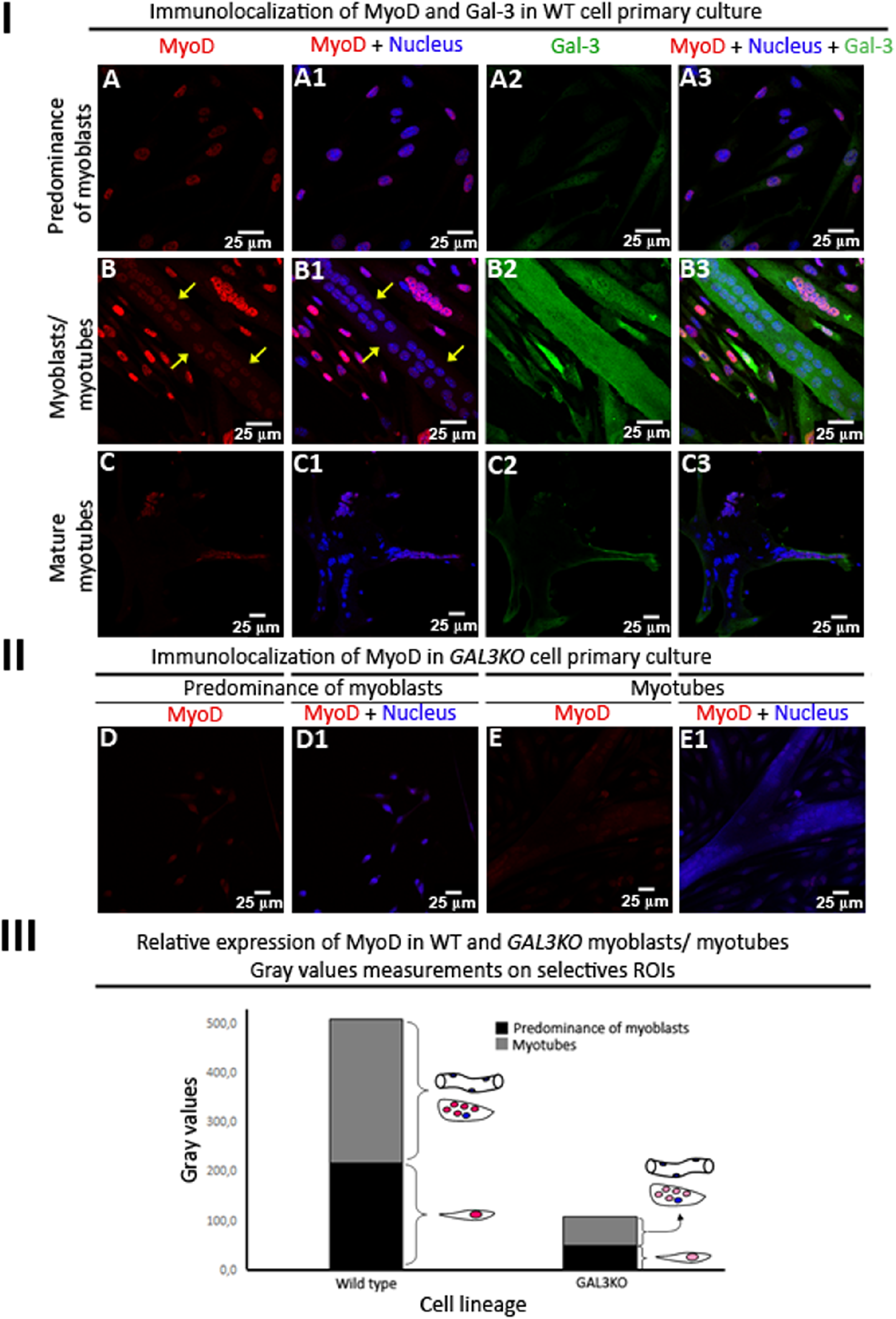
MyoD and Gal-3 are expressed concomitantly during myoblast differentiation. Gal-3 (green), MyoD (red) and nuclei (blue) merged localization during the dynamics of myoblast differentiation to myotubes is shown here. Cells in GM (I-A to I-A3, II-D and II-D1) and in DM (I-B to I-B3, I-C to I-C3, I-E and II-E1). Zero day in I-A to I-A3, II-D and II-D1; 1 day of differentiation in I-B to I-B3, II-E and II-E1; 3 days of differentiation in I-C to I-C3. MyoD starts its expression on day zero (I-A and I-A1), peaking within 1 day of differentiation in proliferative myoblasts and nascent myotubes (I-B and I-B1) but absent on mature myotubes (I-B and I-B1, yellow arrows). After three days of exposition to DM, MyoD (I-C and I-C1) and Gal-3 (I-C2 and I-C3) had their expression drastically reduced on mature myotubes. In *GAL3KO* myoblasts (II-D, II-D1, II-E and II-E1) MyoD presented a very low expression. The bottom scheme (III) represents the relative expression of MyoD during myoblast differentiation; it has a high expression on WT cells, but it is diminished drastically on *GAL3KO* cells. It is important to note that the relative quantity MyoD rate from myoblasts/myotubes remains practically the same (1:1.2).

### Myolesion on *GAL3KO* mice reveal an impaired repair process

The histopathological findings (Figure 1) revealed that *GAL3KO* skeletal muscle responds differently to barium-induced myolesion when compared to WT animals. In Figure 1, the upper-case letters represent a broad view of the muscle tissue while the lower-case letters, a larger magnification under the same conditions. WT mice showed degenerate areas in muscle tissue after one day (Figure 1A, central region and 1a), a similar pattern could be noted in *GAL3KO* mice (Figure 1E and 1e). After 3 days post-injury, the normal muscle presented intense inflammatory infiltrate (yellow arrowhead on the Figure 1B and 1b) and degenerated myofibers, whereas *GAL3KO* presented lower inflammatory infiltrate (yellow arrowhead on the Figure 1F and 1f). At 7 days post-injury, normal muscles are in the resolution phase of tissue repair (green arrows on the Figure 1C and 1c) characterized by basophilic staining and central myonuclei. Intriguingly, *GAL3KO* muscles at 7 days post-injury showed persistent inflammatory infiltrates (yellow arrowheads on the Figure 1G and 1g) along with a structure indicative of fibrosis (bundles in pink tones surrounding the still injured tissue) together with few myofibers in the final state of the repair resolution (green arrows on the Figure 1G). After 14 days, WT mice presented typical morphology of intact muscle (Figure 1D and 1d). However, *GAL3KO* muscle (Figure 1H) exhibited intense inflammatory infiltrate (yellow arrowhead on the Figure 1H and 1h) and myofibers in the final state of tissue repair resolution (Figure 1H and 1h, green arrows) plus fibrotic tissue (Figure 1h, lower right portion).

### *GAL3KO* repairing muscle maintains high osteopontin expression after myolesion

One of the key players in muscle regeneration, osteopontin (OPN) (Pagel et al., 2014) was upregulated after myolesion in *GAL3KO* animals (Figure 2). The localization pattern of OPN-after 1 and 3 days post-myolesion is similar in WT (Figure 2B and 2C, respectively) and *GAL3KO* mice (Figure 2G and 2H, respectively), corresponding to regions of muscular degeneration. At 7 and 14 days, OPN in WT mice is restricted around myofibers (Figures 2D and 2E, respectively). However, in *GAL3KO* animals at the same period, OPN still presented in various muscle regions and around myofibers (Figure 2I and 2J, respectively). The high concentration of OPN in *GAL3KO* mice corroborate with the presence of fibrous scar tissue.

### *GAL3KO* repairing muscle presents the formation of fibrous scar tissue

In order to confirm fibrosis, the sections embedded in paraffin were stained to show collagen fibers in two staining: Sirius Red (Figure 3A and 3C) and Masson’s trichrome (Figure 3B and 3D). Micrographs of 14 days post myolesion were highlighted here. For Sirius Red staining, the myofibers were visualized in light pinkish tone, nuclei in dark blue and collagen fibers in deep red. Under the same condition, Masson’s trichrome staining also revealed light-blue areas for collagen and the remaining myofibers in intense purple staining. In summary, the results obtained by both staining showed extensive areas of fibrosis in muscles of *GAL3KO* animals (Figure 3C and 3D) when compared to their respective controls (WT) after 14 days of barium-induced myolesion (Figure 3A and 3B, respectively).

### The skeletal muscle repair requires transient expression of Gal-3

In order to analyze the participation of Gal-3 in muscle repair process, barium-injured muscular tissue showed the presence of one immunoreactive Gal-3 band at 30 kDa by western blot (Figure 4, I-left panel). Gal-3 demonstrated a differential expression pattern, initially after 1 day, increasing its expression in 2 days and culminating after 3 days. At 7 days post-myolesion Gal-3 expression was diminished and after 14 days its expression was not detected. The second band at 53kDa was identified as desmin, an internal constitutive muscle marker. As an experimental control, the same amount of saline solution was injected (Figure 4, I-right panel), which indicated labeling for desmin only, confirming that Gal-3 was expressed only in the presence of muscle injury.

### Gal-3 presents a dynamic and pleiotropic localization in repairing muscle

During the tissue repair process, as expected, Gal-3 did not exhibit detectable labeling in uninjured tissues (day 0, Figure 4-II-A). After 3 and 7 days (Figure 4-II-B and 4-II-C, respectively), Gal-3 showed to be distributed in multiple sites throughout the injured area, as described next: at 3 days, the lectin was presented in regions with a suggestive pattern of the contour of sarcoplasm membranes (Figure 4-II-B, arrows), with regions external of intact myofibers (Figure 4-II-B, inset and asterisk), seen by dystrophin visualization. At 7 days it was observed in a location restricted to the sarcoplasm (Figure 4-II-C, arrow), which was confirmed by the overlap with β1-integrin (Figure 4-II-C, inset-arrow) and myonuclei (Figure 4-II-C, arrowhead and inset-arrowhead). After 14 days upon completion of regeneration (Figure 4-II-D), Gal-3 was minimally expressed, except for its occurrence in small regions between myofibers.

### MyoD and Myogenin have a decreased expression in the absence of Gal-3

In *GAL3KO* mice, MyoD and Myogenin expression were decreased (Figure 4, III-right panel) when compared with the WT mice (Figure 4-III-left panel), indicating a crucial importance of Gal-3 in these transcription factors expression.

### The formation of myotubes from isolated myoblasts requires a dynamic expression and localization of Gal-3

The differentiation process from myoblasts to myotubes focusing on Gal-3 localization is showed in Figure 5. This lectin was seen inside (Figure 5A, 5B, 5C, 5E and 5F) and outside the cells (Figure 5D). The isolated myoblasts presented variable levels of Gal-3 expression: (i) undetectable (Figure 5A and 5B, yellow arrows), (ii) growing level of expression when myoblasts started to align (Figure 5B, yellow arrowhead) and (iii) high levels in myoblast with a fusiform morphology (Figure 5A, white arrowhead). At the early myotube formation (Figure 5C), Gal-3 was located throughout the cytoplasm (Figure 5C) and myonuclei (highlighted in Figure 5C, arrowhead and inset). As differentiation of myotubes proceeds, we observed that myonuclei showed an aligned fashion (Figure 5C to 5F) together with a redistribution of Gal-3 to cell membrane regions (Figure 5E, white arrow). Interestingly, in non-permeabilized myoblast cells, Gal-3 was located in clusters on the surface of the syncytium (Figure 5D). It is important to highlight that we performed a serum analysis to confirm the absence of Gal-3 on GM and DM culture medium used in all *in vitro* assays (Supplementary Figure S1-A).

The relative expression of Gal-3 during the dynamics of myoblast to myotube differentiation is shown in Figure 5-II. Myoblasts initiate the expression of Gal-3 as they differentiate into myotubes, with an increase of approximately five-fold in its expression, followed by its decreasing.

### The transient expression of MyoD follows the dynamics of Gal-3 expression

To corroborate the previously obtained data, the myoblast/myotube primary culture derived from WT mice was subjected to a triple labeling with Gal-3 (green), MyoD (red) and Nucleus (blue) in Figure 6-I. In fields with a predominance of myoblasts (Figure 6-I-A, 6-I-A1, 6-I-A2 and 6-I-A3) the expression of MyoD is almost concomitant with that of Gal-3. In those fields where the presence of aligned myoblasts and initial myotubes (Figure 6-I-B, 6-I-B1, 6-I-B2 and 6-I-B3), MyoD is highly expressed in aligned myoblasts and nascent myotubes, and has a drastically decreased expression in mature myotubes (Figure 6-I-B and 6-I-B1, yellow arrows). Other micrographs exhibited very low expression of MyoD and Gal-3 in a mature myotube (Figure 6-I-C, 6-I-C1, 6-I-C2 and 6-I-C3).

### *GAL3KO* myoblast have reduced MyoD expression

For a more detailed analysis of the Gal-3 function in muscle repair, we used Gal-3 knockout mice as a source of myoblasts. Prior to cell culture, we confirmed, via western blot, that the fetal bovine and horse serum used in the experiment did not present Gal-3 in its composition (Supplementary Figure S1-B). Our results indicated very low expression of MyoD, both in fields with predominance of myoblasts (Figure 6-II-D, 6-II-D1) and in regions of mature myotubes (Figure 6-II-E and 6-II-E1). For MyoD, a relative quantification indicated that MyoD detection in WT animals is approximately 5x greater than in myogenic cells from *GAL3KO* mice (Figure 6-III); however, the ratio of MyoD present in myoblasts and myotubes remained at the same proportion (approx. 1: 1.2).

### *GAL3KO* myoblasts have Pax7 expression retained even after differentiation

Another interesting finding concerned the expression of the transcription factor Pax7, typical of quiescent satellite cells (Olguin et al., 2007; Shi and Garry, 2006). Pax7 was increased in *GAL3KO* myoblasts (Figure 7), suggesting a possible delay in the conversion of satellite cells to proliferative myoblasts. Note that our western blot for Gal-3 (Figure 7) revealed little fluctuation in its expression during the differentiation process, as the primary cultures were heterogeneous with myoblasts and myotubes.

**Figure 7.**
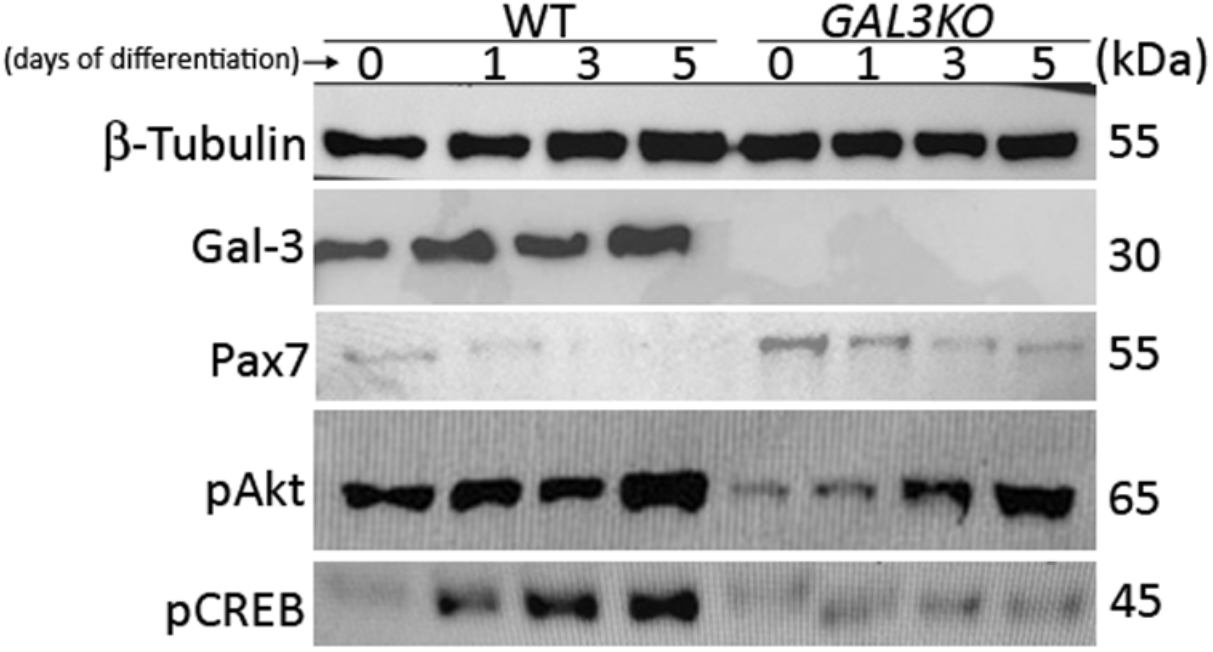
Cell signaling effectors in myoblast to myotube differentiation was decreased, while Pax7 retains its expression. In WT myoblast primary cell culture, Gal-3 is expressed during all the differentiation process together with a decrease of 55 kDa Pax7 expression and increase substantially on the phosphorylation cascade panel for 65 kDa pAkt and 45 kDa pCREB. In *GAL3KO* lineage, Pax7 presents a persistent expression during myoblast differentiation while the phospho cascate, here seen by pAkt and pCREB, were drasticly reduced. β-tubulin is used as standard constitutive protein expression.

### The tyrosine-phosphorylated cascade signaling on key proteins are impaired on *GAL3KO* myoblast cell culture

Analysis of WT myoblasts for phosphorylated proteins demonstrated immunoreactive bands before and after differentiation had initiated (Figure 7). The signaling-targeted proteins Akt and CREB were relatively less in expression in *GAL3KO* cells. The most pronounced result was exhibited by the downstream-phosphorylated nuclear effector pCREB, which appears to be almost missing from *GAL3KO* cells. These data indicate that cell signaling is impaired on *GAL3KO* myoblasts.

### Supplementary data reinforce Gal-3 participation in MRP

The absence of Gal-3 expression in *GAL3KO* was confirmed by immunoblotting 3 days post-myolesion (Supplementary Figure S1-A). On the other hand, only *mdx* mice were positive for Gal-3 expression when compared with wt mice (Supplementary Figure S2-C). Also, Gal-3 localization in muscle tissue (from 4 week old mice) was observed in sarcolemma, myonuclei, and sarcoplasm (Figure S2-E and S2-E-inset)-the same locations observed after acute myolesion in WT mouse (Figure 4-II-B and C). Besides, Gal-3 expression was persistent between 4 to 10 weeks old *mdx* mice (Supplementary Figure S2-F).

## DISCUSSION

Our results indicate that Gal-3 has an essential role in the skeletal muscle repair process after acute myolesion by injection of barium chloride *in vivo* (Caldwell et al., 1990; Doroshow et al., 1985) and in myogenic programming in mouse primary myoblasts cells *in vitro*. The histopathological analysis showed that the tissue repair dynamics of *GAL3KO* skeletal muscle is negatively affected when compared to WT animals, remaining in the resolution phase until the 14^th^ day post-myotrauma. The impaired tissue repair is multifactorial and, in these mice, may be associated with the delay in the recruitment of inflammatory cells to the site of myolesion, which usually occurs with an accumulation at 3 days post-injury. In *GAL3KO* animals, inflammatory cells were observed and analyzed throughout the post-injury periods (3-14 days), suggesting that Gal-3 plays an important role in chemotaxis and secretion of immune mediators leading to deficiency in the cellular homeostasis. Gal-3 has an inflammatory role in several immunological processes such as neutrophil extravasation, recognition, and phagocytic removal, all of which are intrinsically correlated with MRP (Loimaranta et al., 2018; Nieminen et al., 2005; Sano et al., 2003). Also, Gal-3 induces macrophage polarization to the M2 phenotype that can downregulate inflammation, limit tissue injury, and/or promote muscle repair (MacKinnon et al., 2008). The orchestrated production of pro-or anti-inflammatory cytokines by macrophages M1/M2 and infiltrating inflammatory cells, such as TNF-α, IFN-γ, IL-6, and IL-10 is crucial for MRP (Tidball et al., 2014). In regenerating muscle, IL-6 has been described as a pivotal cytokine produced by infiltrating macrophages, neutrophils, fibro/adipogenic progenitors (Joe et al., 2010), and satellite cells (Muñoz-Cánoves et al., 2013). It has been demonstrated *in vitro* that human myoblasts produce IL-6 in response to TNF-α and IL-1β, and the increased levels of this cytokine contribute to stimulate satellite cells to proliferate, differentiate, and fuse (Gallucci et al., 1998; Muñoz-Cánoves et al., 2013). In addition, IL-6 knockout mice are not able to respond properly to tissue injury as a result of loss in cellular response (Fattori et al., 1994). In this context, Gal-3 may contribute to the inflammatory status of the damaged tissue by inducing leukocytes chemotaxis and its activation and promoting IL-6 release from tissue fibroblasts (Filer et al., 2009). Thus, the absence of Gal-3 may have a functional impact that implies in the deregulation of immune cells activation, as well as the release of cytokines and chemokines, favoring Gal-3 as a key part of MRP.

During the onset of MRP, a transient extracellular matrix appears in the area of inflammatory infiltration (Serrano et al., 2011), which is solved after completion of the tissue repair. If a high number of myofibroblasts are activated while the matrix is being solved, the fibrotic elements start to replenish the injured tissue, playing and important role in the development of fibrotic diseases (Hinz et al., 2007; Kis et al., 2011; Rosenbloom et al., 2017). Several studies has described Gal-3 as a fibrotic marker that represents a worse prognostic for heart failure (Tanaka et al., 2020). Besides, this lectin contributes positively to atrial fibrosis by the activation of the TGF-β1/α-SMA/Col I pathway (Shen et al., 2018) and upregulates the activation of myofibroblasts that culminate in hepatic fibrosis (Henderson et al., 2006). On myocardial macrophages, increased Gal-3 expression and its binders on matrix extracellular and fibroblast, accelerate cardiac fibroblasts proliferation and collagen production (Sharma et al., 2004). On the other hand, there is evidence that Gal-3 expression is correlated with less fibrosis in lung tissue in chronic air inflammation (López et al., 2006) and experimental diabetic glomerulopathy (Pugliese et al., 2001). *In vivo*, the treatment with Gal-3 can modulate eosinophil and T cell infiltration through IL-5 downregulation (Del Pozo et al., 2002) and consequently reducing collagen deposition in the lung, improving airway remodeling (López et al., 2006). In diabetic glomerulopathy, it has been observed that the presence of Gal-3 also acts as advanced glycation end products (AGEs) receptor, which is similar to macrophage scavenger function. When these AGEs products are removed from tissue injury, the cell homeostasis and remodeling processes are favored. In agreement with this literature, histopathological analysis of *GAL3KO* mice showed a structure confirmed by specific staining for collagen fibers, indicating persistent fibrous scar tissue up to 14 days after myolesion. Thus, the lack of Gal-3 is associated with induction of scar formation of fibrotic tissue in the repairing skeletal muscle, which suggests that this lectin can act improving the homeostasis along this process and protecting the tissue from severe injury.

Interestingly, our results indicate that during the early stages of the MRP, osteopontin (OPN) expression and release occur in both WT and *GAL3KO* mice. However, in the final stages, OPN expression diverge: in WT animals, it is decreased and its location is restricted to the myofiber environment, and in *GAL3KO* animals OPN expression is still high, located in regions indicative of fibrous tissue and around myofibers. OPN has been described a key protein for muscle repair by Hirata *et al*. (2003) with an increase of 118.5x after acutely induced myolesion in mice and with high levels of OPN mRNA (145.6x) in chronic muscular lesion in *mdx* strain by Porter *et al*. (2002). OPN promotes adhesion, proliferation, migration, and chemotaxis, among other processes; all of which regulate physiological and pathological situations such as tissue repair, inflammation and fibrosis (Hirata et al., 2003; Pagel et al., 2014; Porter et al., 2002). In the specific case of myoblasts, OPN promotes their multiplication and it is required for the final maturation of the newly synthesized myofibers (Pagel et al., 2014). Taken together, these data suggest that the absence of Gal-3 disrupts the OPN negative turnover, which would impair muscle tissue homeostasis.

Additionally, we observed increased Gal-3 expression in proliferating myoblasts from primary cell culture during myogenesis. The role of Gal-3 on the efficiency of myogenesis was described recently in C2C12 cells (Rancourt et al., 2018) and our findings corroborate the findings regarding overall processes of myoblast to myotube differentiation. Nevertheless, the morphology of myotubes becomes much more evident in primary cell culture and the use of these kinds of cell culture allowed us to obtain data at a closer physiological level than C2C12 cells approach would permit (Abdelmoez et al., 2020).

Gal-3 is translated in the cytoplasm as a soluble protein and exported outside the myoblasts, then may link itself to glycomolecules at the myoblast cell surface in a scattered and diffuse fashion. The aligned myoblasts, the nascent myotubes and the early myotubes exhibited very high Gal-3 expression. As fusion proceeds, however, Gal-3 is rearranged at the cell surface level on a clustered pattern, likely contributing to activation of a signaling cascade. Additionally, inside the nascent myotubes the expression of Gal-3 is increased abruptly up to 5x that coincides with a nucleocytoplasmic location. Interestingly, we obtained the same subcellular location for Gal-3 in both *in vivo* and *in vitro* experiments: Gal-3 was also strongly labeled in the sarcolemma for newly regenerated myofibers and in the final myotubes maturation in cell culture. It has been reported that Gal-3 occurs in a basolateral membrane location in mouse enterocytes (Delacour et al., 2008). As myotubes mature, the expression of Gal-3 decreases abruptly (similar data was obtained by (Rancourt et al., 2018), and remains linked only in regions of the sarcolemma, where it likely assists the stabilization of the myotube membrane. Rancourt and colleagues (2018) further suggested that Gal-3 drives cellular fusion processes rather than stages of cell differentiation *per se*. Our results are consistent, in part, with these previous findings, since the amount of myogenic transcription factor MyoD is much lower in *GAL3KO* myoblasts culture. However, the ratio between MyoD+ nuclei observed in mononuclear cells (myoblasts) and syncytia (myotubes) remains similar in both myoblast lineages. These results suggest a relevant participation of Gal-3 in the transient assembly of the newly formed membrane by the fusion of myoblasts (a graphical scheme of MyoD and Gal-3 co-expression during myoblast differentiation is provide in Supplementary Figure S3). It is tempting to infer that this location of Gal-3 in the sarcolemma could create a specific microenvironment, influencing biophysical properties of the cytoplasmic membrane, resulting in the stabilization or enhancement of myogenic cell membranes designed to function under constant mechanical stress. The positive detection of Gal-3 in myonuclei may implicate its participation on spliceosome machinery and *snRNP* complexes, which could control cell cycle and prevent cellular programmed cell death (Haudek et al., 2010; Yang et al., 1996). As reported by Haudek and colleagues (2010), Gal-3 exhibits nucleocytoplasmic shuttling, or repeated movement in a bidirectional way across the nuclear pore complex, which has been observed in several cell types.

Moreover, *GAL3KO* myoblasts expressed a greater amount of the Pax7 transcription factor, typical of quiescent satellite cells, even during the later differentiation process, suggesting that there is a delay in the programming of muscle differentiation comparing with WT mice. Normally, Pax 7 expression on satellite cells decreases whilst the MyoD increases culminating with myoblast activation, proliferation, fusion and differentiation (Sambasivan et al., 2011). Indeed, the MRP process requires a sequence of phosphorylation events, including Akt and cAMP response element binding protein (CREB) residues, that are linked to promote MyoD expression and consequently myoblast differentiation (Stewart et al., 2011; Zhu et al., 2017). Others have shown that pCREB interacts with MyoD (Magenta et al., 2003) and CREB with Akt (Du and Montminy, 1998). In this study, *GAL3KO* myoblasts was found decreased levels of tyrosine phosphorylated on residues of Akt and CREB transcription factors, reflecting the deficiency of resolution of the injured muscle.

Our results also corroborate studies of others demonstrating that both MyoD and pCREB are diminished in the absence of Gal-3, which may delay myoblast differentiation. Recently, Zhang and colleagues (2018) showed that the treatment of pulmonary artery endothelial cells *in vitro* with Gal-3 increases MyoD expression, which can induce an alpha-smooth muscle actin (α-SMA)-positive phenotype through the transdifferentiation of these cells into vascular smooth muscle cells, thus indicating a positive feedback between Gal-3 signaling and MyoD expression (Zhang et al., 2018). Another group of proteins also appear to play a crucial role in skeletal muscle inflammation and fibrosis: Matrix Metalloproteinases (MMPs) and Tissue Inhibitors of Metalloproteinases (TIMPs), as reviewed recently (Alameddine and Morgan, 2016). It is known that MMPs are essential to myoblast fusion (Couch and Strittmatter, 1983), and also other functions, such as described for MMP2 that is involved in satellite cell activation, whereas MMP9 contributes to reduces excessive inflammation (Alameddine and Morgan, 2016). Interestingly, Gal-3 was identified as a substrate for MMP2 and MMP9 (Ochieng et al., 1994). Thus, the absence of Gal-3 may disrupt the intrinsic balance between the extracellular matrix and other elements that participate in the homeostasis and muscle remodeling after myolesion.

In conditions of chronic muscular lesions such as seen in *mdx* mouse strain, prior studies reported that the mRNA of Gal-3 was overexpressed ∼91x in comparison to the skeletal muscle of WT ones (Porter et al., 2002). Concerning chronic muscular degeneration, we confirmed Gal-3 expression in *mdx* mice (Supplementary Figure 3), demonstrating that it is expressed during the first 10 weeks of animal lifespan and has a diverse location in the muscle tissue, like an unorganized pattern on the sarcoplasm, into the myonuclei and in the sarcolemma in 4 weeks old animals. Such data suggests that the high expression of Gal-3 in the dystrophic muscles, compared to the WT ones, may contribute to muscle homeostasis.

## CONCLUSION

The data presented here indicate that Gal-3 expression in the inflamed skeletal muscle is initiated upon the induction of lesion and becomes undetectable with the muscle repair. In addition, Gal-3 expression is associated with altered expression levels of transcription factors involved with myoblasts differentiation to myotubes. These results suggest that endogenous Gal-3 plays a critical role in skeletal muscle homeostasis upon myolesion, as ilustrated in Supplementary Figure S4.

## LEGEND TO FIGURES

**Supplementary Figure S1.**
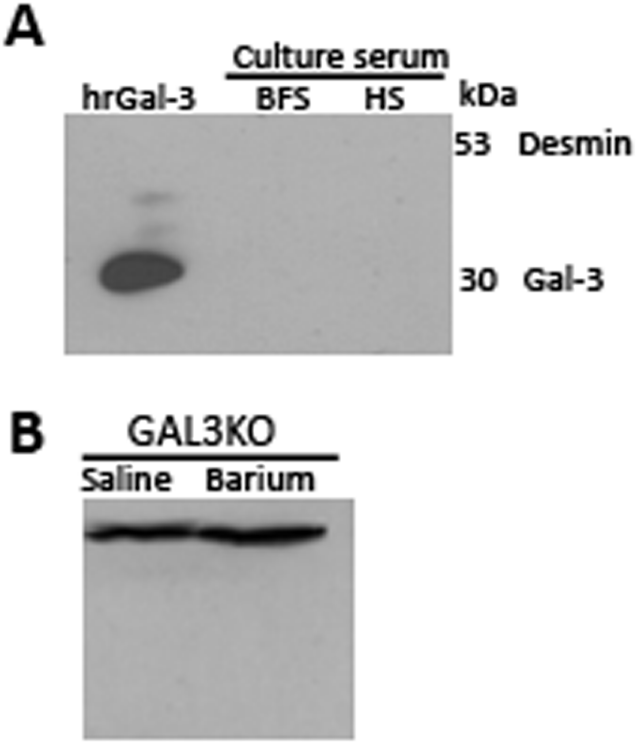
SDS-PAGE proved that the serum used in this study contains no Gal-3 (A) and the *GAL3KO* murine muscle does not express this lectin (B), even after 3 days post-injury.

**Supplementary Figure S2.**
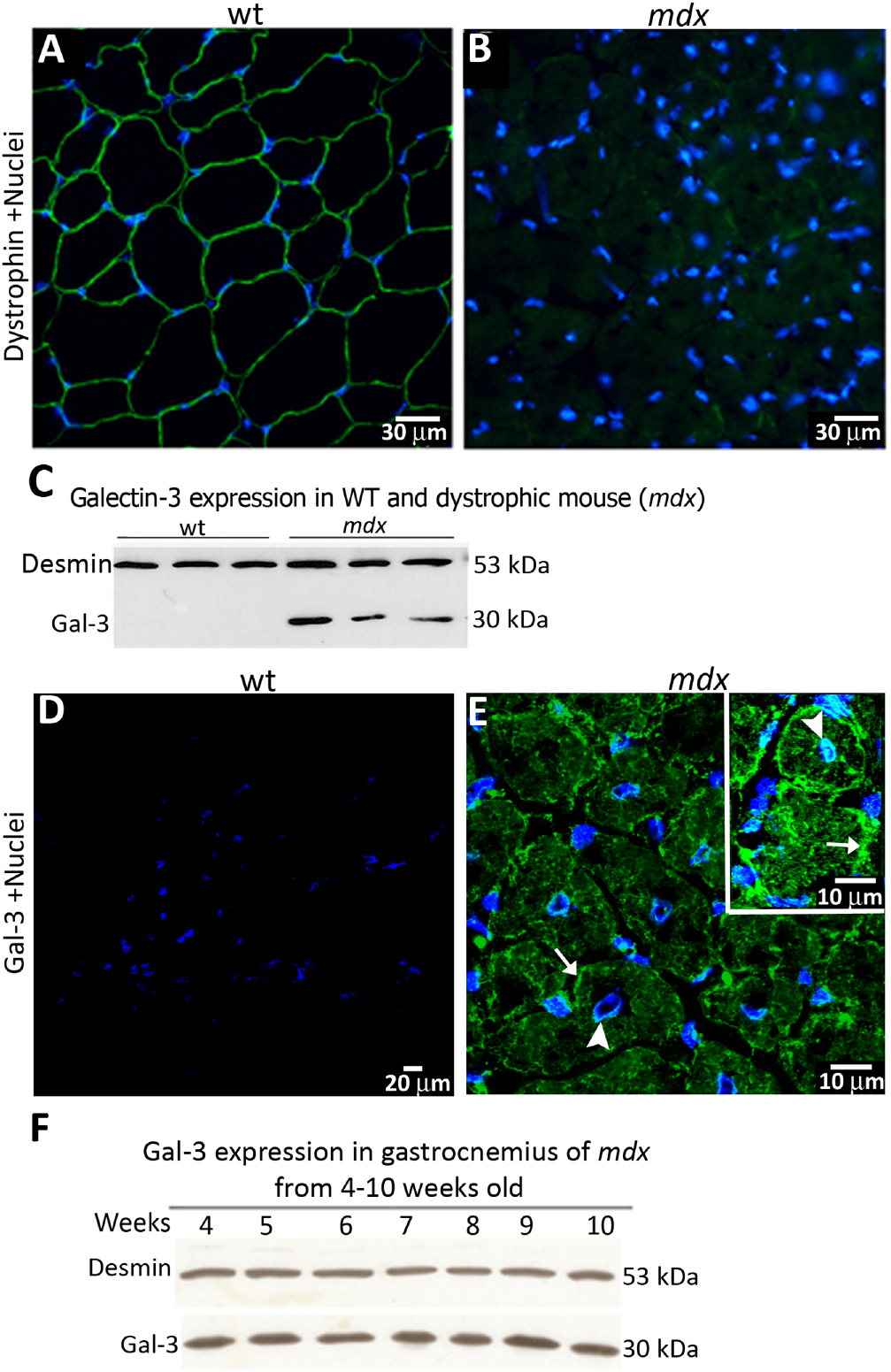
Gal-3 is constitutively expressed in the dystrophic mouse (*mdx*). Gal-3 is constitutively expressed in the dystrophic mouse (*mdx)*. Immunofluorescence to dystrophin (A, wt-in green) confirms *mdx* phenotype (B, *mdx*). Three *mdx* mice were subjected to WB analyses, showing that only *mdx* expresses Gal-3 (C); Desmin were used as constitutive muscle marker (C). Immunofluorescence images from Gal-3 showed that it is not detectable on wt mice (D), but is located on myonuclei (E, arrowhead and E, inset-arrowhead), sarcoplasm and sarcolemma (E, arrow and E, inset-arrow). Besides, *mdx* mice express Gal-3 during all the analyzed period-from 4 to 10 weeks-old (F).

**Supplementary Figure S3.**
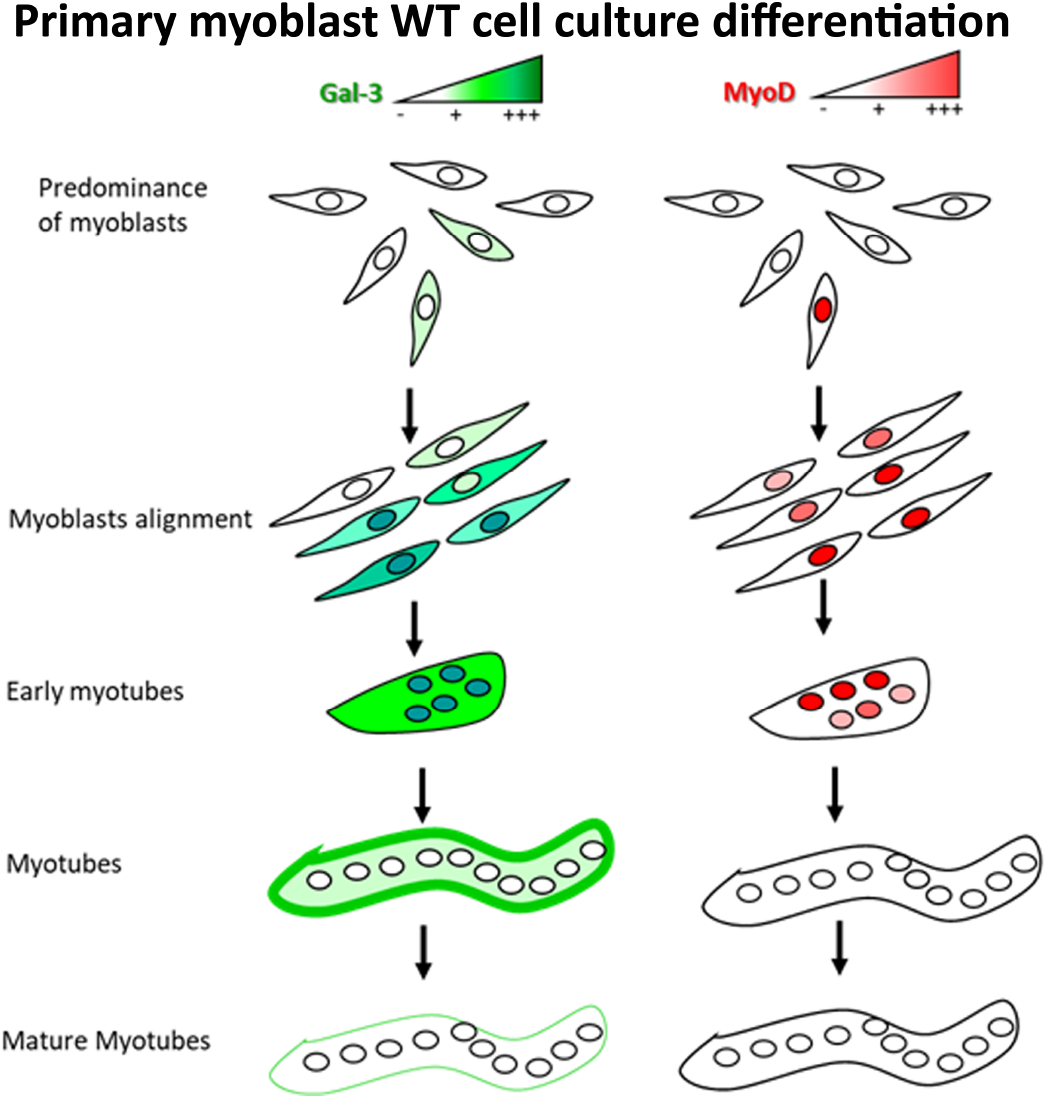
Scheme of Gal-3 and MyoD co-expression during myoblast differentiation to myotubes.

**Supplementary Figure S4.**
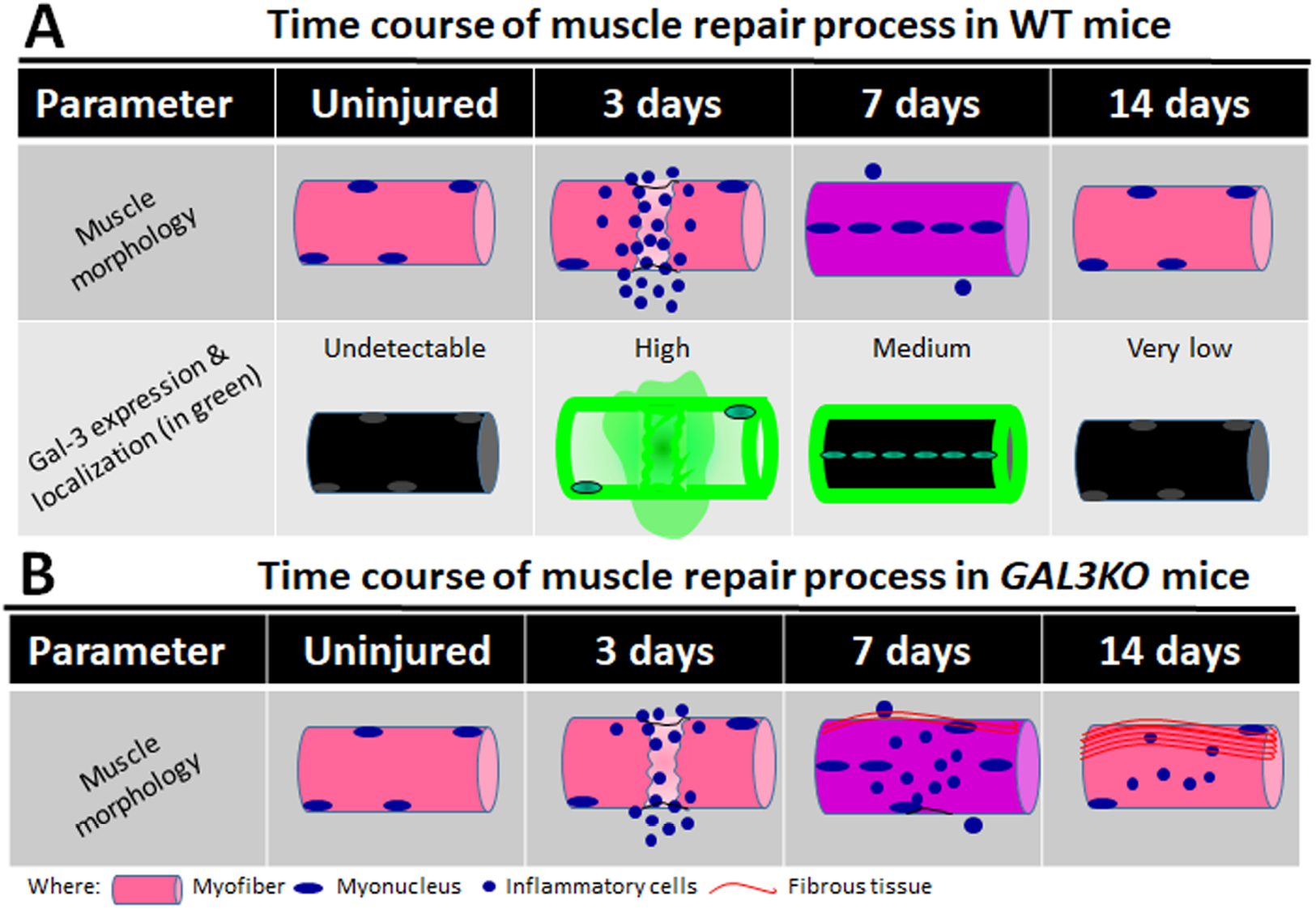
Overview of myolesion repair in WT (A) and *GAL3KO* (B) mice lineages. Note that the absence of Gal-3 impairs the MRP.

## Data Availability

The data used to support the findings of this study are included within the article and supplementary data.

